# Activation of Smad 2/3 signaling by low shear stress mediates artery inward remodeling

**DOI:** 10.1101/691980

**Authors:** Elizabeth Min, Nicolas Baeyens, Rui Hu, Zhenwu Zhuang, Minghao Chen, Billy Huang, Georgia Zarkada, Angela Acheampong, Kathleen McEntee, Michael Simons, Anne Eichmann, Martin A. Schwartz

## Abstract

**Rationale:** Blood vessel remodeling in response to changes in tissue demand is an important aspect of fitness and is often compromised in vascular disease. Endothelial cell (EC) sensing of fluid shear stress (FSS) governs vessel remodeling to maintain FSS at a specific magnitude or set point in healthy vessels.

**Objective:** The purpose of this study was to understand how shear stress-regulated Smad 2/3 contributes to artery remodeling.

**Methods and Results:** We found that shear stress induces Smad 2/3 phosphorylation, nuclear translocation, and gene expression in ECs. Nuclear translocation and gene expression are maximal at low and decrease at high FSS. Reducing flow in the mouse carotid by ligation of branch vessels induces Smad2 nuclear localization in vivo. Activation of Smad 2/3 by FSS requires the Type I TGFβ family receptor Alk5 and the transmembrane protein Neuropilin-1. Flow activation of Smad 2/3 is mediated by increased sensitivity to BMP9 but not BMP10 or TGFβ. By contrast, flow activation of Smad 1/5 is maximal at physiological FSS and requires BMP9 or 10 binding to Alk1 and Endoglin. EC-specific deletion of Alk5 in mice blocks low flow-induced inward remodeling after carotid ligation.

**Conclusions:** Together, these data elucidate a novel pathway that mediates low flow-induced inward artery remodeling. These results may be relevant to inward remodeling in diseased vessels where Smad 2/3 is activated by pathological stimuli.

## Introduction

The vertebrate vasculature system is designed to adjust local perfusion to match metabolic demands, thus optimizing tissue function and ultimately survival. For example, hypoxia induces secretion of factors that dilate small resistance vessels on short time scales, and, on longer time scales, angiogenic factors that increase vessel density [1]. Both of these adjustments increase flow through the larger arteries that feed the affected tissue, inducing their outward remodeling. Conversely, disuse or atrophy of a tissue results in decreased vessel density and inward remodeling of the feeder arteries [2].

Endothelial cells (ECs) that line the inner surfaces of blood vessels sense and transduce signals from hemodynamic fluid shear stress (FSS) to regulate vessel remodeling [3]. Multiple studies have shown that surgical interventions to increase or decrease flow through an artery stimulate outward or inward remodeling, respectively [4, 5]. Current evidence supports a model in which the endothelium encodes a FSS set point that mediates homeostasis. Sustained FSS above or below this value induces outward or inward remodeling, thereby restoring FSS to its original value. By contrast, FSS near the setpoint stabilizes vessels. The set point varies for different types of ECs, corresponding to the physiological level of shear stress appropriate to the specific vessel type. For HUVECs, inflammatory pathways were suppressed and Smad 1/5 maximally activated in the physiological FSS range of 10-30 dynes/cm^2^, suggesting a role for the TGFβ family ligands and receptors in shear stress regulation [6].

Signaling by TGFβ family components regulate a wide variety of cellular responses including proliferation, differentiation, and migration from early developmental stages to adult life as well as in disease [7]. Activation is initiated by homologous ligands that have been grouped into three subfamilies: TGFβs, bone morphogenetic proteins (BMPs), and activins [8]. These proteins bind to Type I and Type II transmembrane receptors with cytoplasmic serine/threonine kinase domains. Ligand binding initiates receptor activation and phosphorylation of receptor-regulated Smads (R-Smads) in their C-termini. Non-kinase Type III receptors sometimes facilitate ligand binding and signaling. Smad C-terminal phosphorylation facilitates their binding to Smad 4 and nuclear entry of this dimeric complex which directly binds target gene promoters to regulate gene expression. As Smads are continuously shuttled in and out of the nucleus, nuclear accumulation correlates with to their activation (i.e., ability to induce target gene transcription) [9, 10].

Our previous work showed that activation and nuclear translocation of Smad 1/5 near the physiological set point was downstream of its receptors Alk1 and Endoglin. Activation by flow was mediated by a ~20-fold increase in sensitivity to the circulating TGFβ family ligands BMP9 or BMP10 [11]. Consistent with a role in vascular stabilization, mutation or deletion of these receptors induces vascular malformations that are fragile and prone to rupture [12]. Additional studies showed that the related Smad 2/3 pathway is also activated by flow but in this case, it was suppressed by physiological laminar FSS and activated by a low/oscillatory FSS profile, typical of atherosclerosis-susceptible regions of arteries [13]. These results prompted us to examine the role of Smad 2/3 signaling in flow-dependent vascular remodeling in more detail.

## Methods

### Cell culture

Primary HUVECs were obtained from the Yale Vascular Biology and Therapeutics core facility. Each batch consists of cells isolated and pooled from three different donors. Cells were cultured in M199 (Gibco) supplemented with 20% FBS, 1X Penicillin-Streptomycin (Gibco; 15140122), 100ug/mL heparin (Sigma), and 150ug/mL ECGS (referred to as complete M199). HUVECs were used between passages 3 and 6 for experiments.

### Shear stress

HUVECs were seeded on tissue culture plastic slides coated with 20μg/mL fibronectin for two hours at 37°C and grown to confluence. For short-term experiments, cells were starved for at least four hours in complete M199 medium diluted 10-fold with serum-free M199 to obtain a final concentration of 2% FBS. Shear stress with a calculated intensity of 1 dyn/cm^2^ to 30 dyn/cm^2^ was applied in parallel flow chambers as described [14] for the indicated times. For shear stress experiments conducted with ligands or blocking antibodies, HUVECs were starved for four hours in serum-free M199 medium containing 0.2% BSA. Recombinant human BMP9 (R&D; 3209-BP-010), BMP10 (R&D; 2926-BP-025), or TGFβ2 (R&D; 302-B2-002) were added in the indicated amounts. BMP9 and BMP10 blocking antibodies [15] were generously provided by Genentech (South San Francisco, CA).

### siRNA transfection

HUVECs were transfected using RNAiMax (Invitrogen; 13778100) in Opti-MEM (Gibco; 31985070) at a final siRNA concentration between 10nM to 15nM according to the manufacturer’s instructions. EGM-2 was replaced with complete M199 12 hours after transfection. Cells were used for experiments three to four days after transfection. ON-TARGET plus Smartpool siRNAs from Dharmacon were used against human Smad 2 (L-003561-00-0005), human Smad 3 (L-020067-00-0005), human Alk5 (L-003929-00-0005), and human Nrp1 (L-019484-00-0005).

### Lentiviral transduction

293Tx cells were cultured in DMEM (Gibco) supplemented with 10% FBS and 1X Penicillin-Streptomycin (Gibco; 15140122). At least 24 hours before transfection, they were transferred to medium without antibiotics. For virus production, cells were transfected with lentiviral plasmids and packaging plasmids using Lipofectamine 2000 (Thermo Fisher Scientific; 11668019) according to the manufacturer’s instructions in Opti-MEM medium (Gibco; 31985070). Supernatants containing lentivirus was collected 36-72 hours after transfection, passed through a 0.22μm filter, and added to HUVECs. Polybrene was added at a final concentration of 8ug/mL to maximize transduction efficiency. HUVECs were incubated with virus for 24 hours and plated for siRNA transfection 3-5 days after transduction.

### Reverse transcription and qPCR

To assay gene expression, HUVECs were washed with PBS and mRNA was extracted using the RNeasy kit (Qiagen; 74106) according to the manufacturer’s instructions. The cDNA was synthesized (Bio-Rad; 170-8890) and used for qPCR analysis (Bio-Rad; 172-5121) using Bio-Rad CFX94 according to manufacturer’s instructions. Primers used for qPCR include human GAPDH (forward 5’ GTCGCTGTTGAAGTCAGAGG 3’ and reverse 5’ GAAACTGTGGCGTGATGG 3’), human Fibronectin (forward 5’ AAACTTGCATCTGGAGGCAAACCC 3’ and reverse 5’ AGCTCTGATCAGCATGGACCACTT 3’), and human Integrin α5 (forward 5’ ATAGGGTGACTTGTGTTTTTAGG 3’ and reverse 5’ AAAGACATGATTGCTAAGGTCC 3’). The mRNA expression for each gene was normalized to the expression of endogenous GAPDH and relative expression level was calculated using the ΔΔCt method. PCR amplification consisted of 5 minutes of an initial denaturation step at 95°C, followed by 35 cycles of PCR at 95°C for 15 seconds and 60°C for 30 seconds.

### Immunofluorescence

HUVECs were washed with PBS, fixed for 10 minutes with 3.7% PFA in PBS, permeabilized for 10 minutes with 1% Triton X-100 in PBS, blocked for 30 min with Starting Block blocking buffer (Thermo Fisher Scientific; 37542) at room temperature, and incubated in primary Smad 2/3 antibody (1:500; Cell Signaling; #8685S) diluted in the Starting Block overnight at 4°C. Cells were washed with PBS, incubated with secondary antibody and DAPI for 2 hours at room temperature, washed again with PBS, and mounted. Images were captured with a 20x objective on a PerkinElmer spinning disk confocal microscope. Image analysis of Smad translocation was performed as described [16].

### Western blotting

HUVECs were washed with PBS and extracted in Laemmli sample buffer. Samples were run on SDS-PAGE gels and transferred onto nitrocellulose membranes. Membranes were blocked with 5% milk in TBST and probed with primary antibodies overnight at 4°C. Primary antibodies used were: Smad 1 (1:1000; Cell Signaling; #9743), Smad 2/3 (1:1000; Cell Signaling; #8685S), P-Smad 1/5/8 (1:1000; Cell Signaling; #13820), P-Smad 2/3 (1:1000; Cell Signaling; #8828), Linker P-Smad 2/3 (1:1000; Cell Signaling; #3104), Alk1 (1:1000; R&D; #AF370), Alk5 (1:500; Abcam; #31013), Nrp1 (1:2000; Cell Signaling; #3725), Endoglin (1:2000; R&D; #AF1097), Actin (1:5000; Santa Cruz; #130656), and Vinculin (1:5000; Sigma; #V9131). HRP-conjugated mouse secondary (Vector Laboratories; #PI-2000) and rabbit secondary (Vector Laboratories; PI-1000) antibodies were used at 1:10,000.

### Animals

8-wk-old male and female C57BL/6J mice (Charles River, Indianapolis, ME) were used in this study. These mice were housed in groups, in a light- (12-h light cycle) and temperature- (69 °F) controlled environment. They were fed a pellet rodent diet, ad libitum, and had free access to water pre-procedure. The study conformed to the guidelines for the Care and Use of Laboratory Animals published by the US National Institutes of Health (NIH Publication No. 85-23, revised 1985) and was approved by the Institutional Animal Care and Use Committee.

### Partial ligation of the right carotid artery

Mice were anesthetized using inhalation of 2% isoflurane through a vaporizer with 100% oxygen. Nair was applied in the neck between the mandible and the sternum to remove the hair. Betadine and 70% ethanol were used to sterilize the surgical area. All surgical manipulations were performed under a dissecting microscope, maintaining aseptic conditions on a heated surgical pad at 37 °C. The cervical skin was cut in the midline. Then the right common carotid artery and its bifurcation were bluntly dissected to expose all four distal branches: external carotid artery, internal carotid artery, occipital artery and superior thyroid artery. The external carotid below the superior thyroid was tied off using 6-0 silk suture. The internal carotid distal to the occipital artery was also tied off using 6-0 silk suture. Thus, only the occipital artery remained patent. The skin was closed with a 6-0 prolene suture. Fully conscious animals were returned to their original cages. Buprenorphine was administered subcutaneously at 0.05 mg/kg, once preemptively, then every 12 h for 72 h. The survival rate for the surgery was 94%.

### Carotid ultrasound examination

To validate whether the partial ligation reduced flow, carotid flow was examined utilizing a commercially available system (the Vevo 770, Fujifilm, Visualsonics Inc) one day post-ligation. Mice were positioned on an imaging stage with a heating system to maintain physiological body temperatures during imaging. Electrocardiographic leads were attached to each limb. Pulse wave Doppler was also recorded in a para-trachea long-axis views in both the right and left common carotid. Images from these studies were recorded digitally for subsequent analysis. Successful partial carotid ligation was indicated by overall reduction in flow (~80-90%) in the right common carotid compared with the left side, with reversal of flow towards aortic inlet during diastole.

### Immunohistology

Carotid arteries were harvested after euthanasia. Briefly, animals were first perfused with PBS then with a fixative solution (4% paraformaldehyde) by injection through the cardiac left ventricle. Arteries were then removed, further fixed in 4% PFA for 2 hours and embedded in paraffin. Thin transverse sections (5 micrometers) were cut approximatively 5 mm upstream of the ligation site. Sections were then de-parafinized and rehydrated. Antigens were recovered by immersion for 20 minutes in an EDTA buffer at pH 8 at 98°C. Slides were then blocked with SuperBlock TBS buffer (Thermoscientific) for 1 hour. Slides were incubated overnight at 4°C with primary Phospho-Smad 2 (ThermoFisher; #4-244G) antibody diluted 1:200 in blocking buffer. Samples were washed 4X then incubated with secondary antibody (anti-rabbit AlexaFluor 647, Molecular probes, Thermoscientific). Slides were then mounted in FluoromountG (Southern Biotech).

## Results

### Smad 2/3 activation by shear stress

We first exposed HUVECs to 12 hours of LSS ranging from 1 to 30 dynes/cm^2^ and quantified Smad 2/3 nuclear translocation. Nuclear Smad 2/3 staining was maximal under low FSS (1-5 dynes/cm^2^) and decreased under higher FSS, including shear in the physiological range (Figure 1A-B). Parallel assessment of cell alignment showed that ECs aligned mainly at physiological LSS as expected (Figure 1C), confirming the normal dose response to FSS. To test the functional significance of Smad 2/3 nuclear localization, we examined two genes, fibronectin and Integrin α5, that were recognized as direct Smad 2/3 target genes in a CHIP assay with antibody to Smad 2 (data not shown). Both fibronectin and Integrin α5 mRNA levels in HUVECs peaked at low shear and depleting Smad 2/3 with siRNA knockdown blunted their expression (Figure 1D-F).

**Figure 1:**
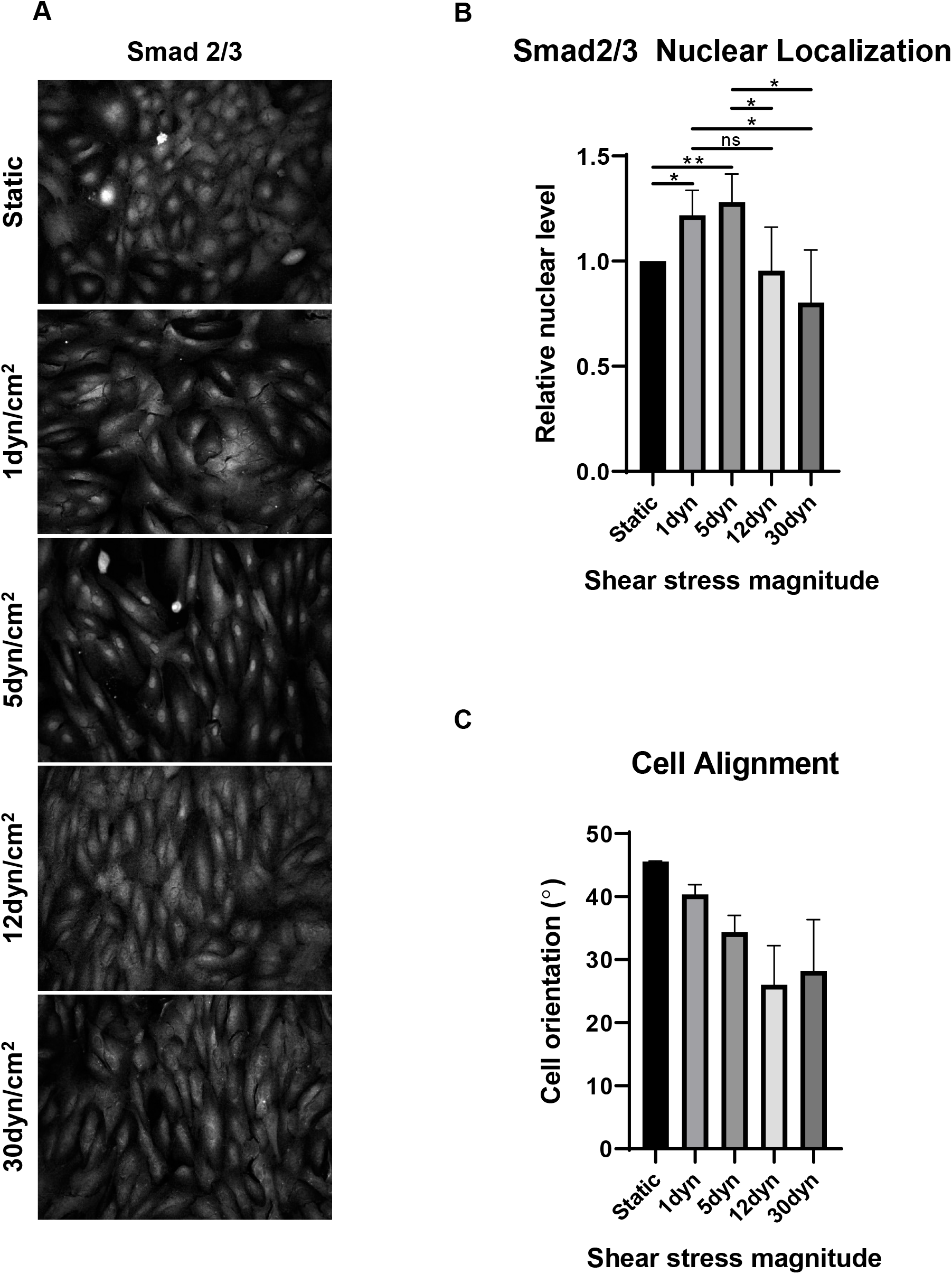
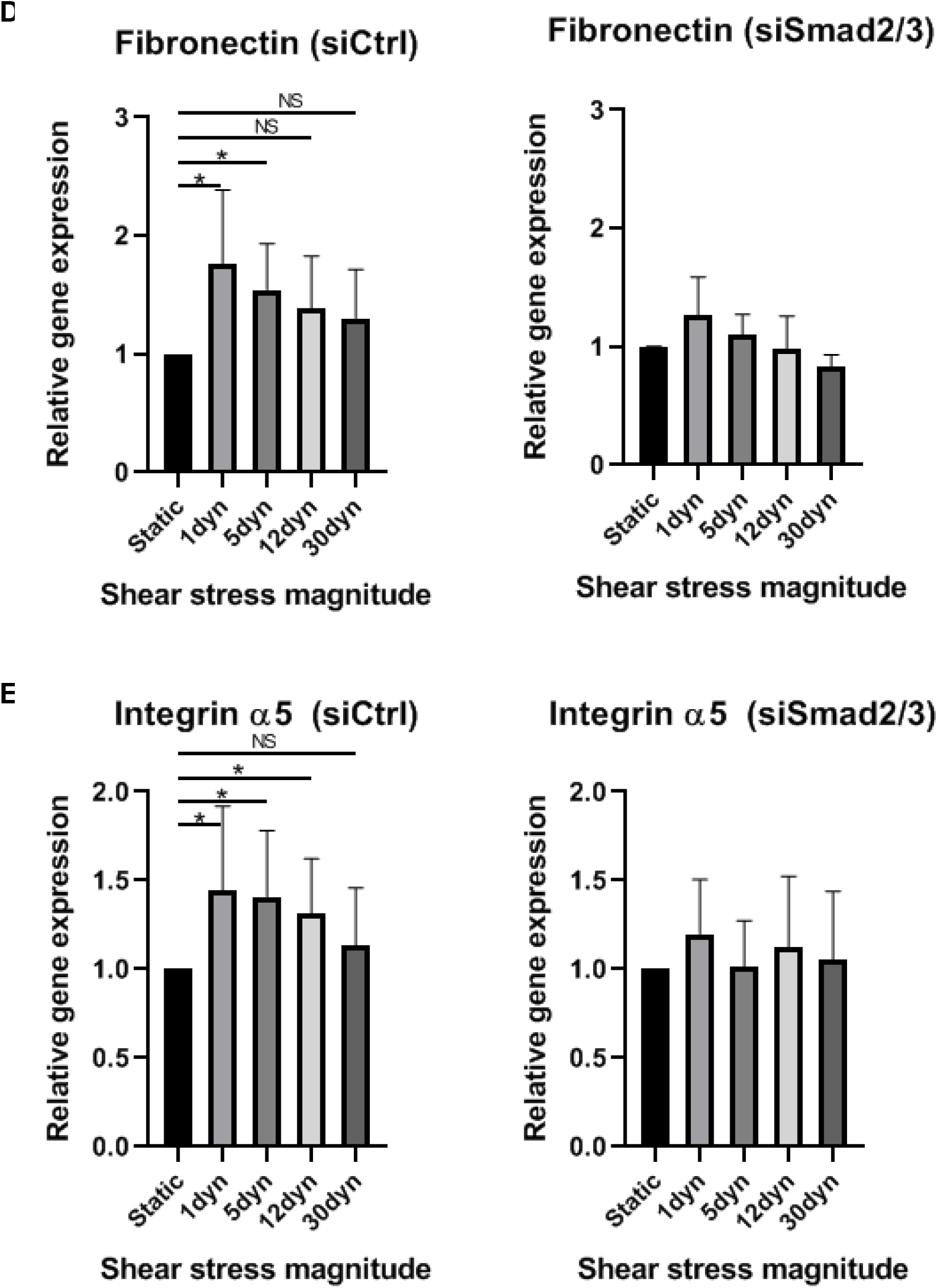
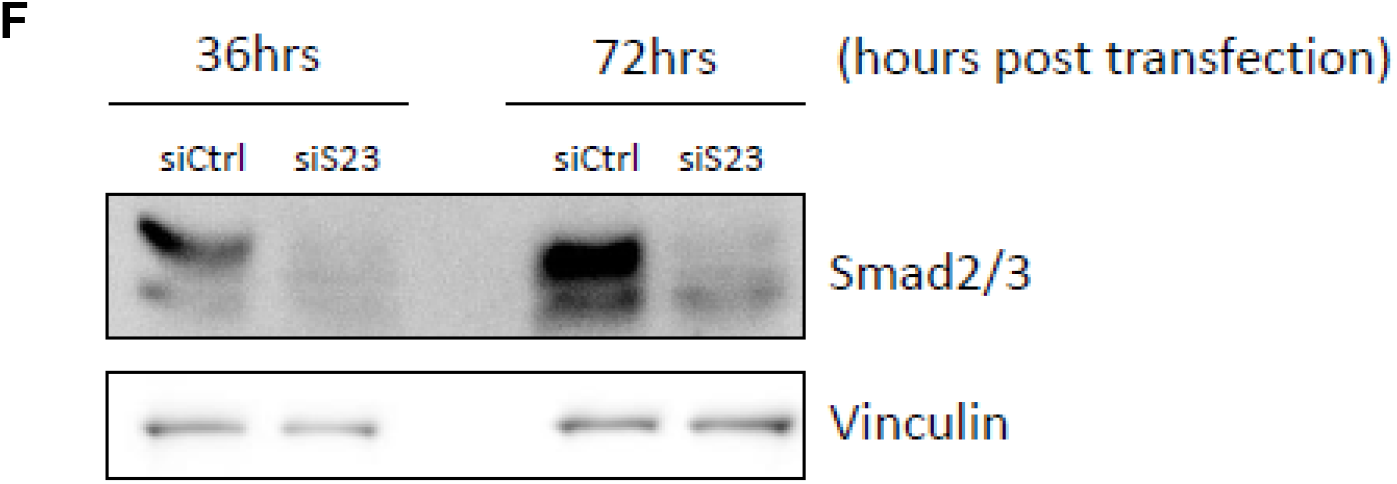
Smad 2/3 regulation by shear stress. **A:** HUVECs were exposed to LSS for 12 hours, fixed, and stained for Smad 2/3. **B:** Quantification of Smad 2/3 nuclear localization. Values are means ± SEM, n=4. **C:** Cell alignment was quantified by measuring the angle between the direction of flow and the long axis of the cell. Values are means ± SEM, n=4. **D-E**: Fibronectin and Integrin α5 mRNA levels were assessed by qPCR. Values are means ± SEM, n=4. **F:** HUVECs were transfected with Smad 2 and Smad 3 siRNA. At the indicated times, cells were harvested and Smad2/3 expression assessed by Western blotting. ^*^ p<0.05, ^**^ p<0.01

### Smad 2/3 phosphorylation

The TGFβ receptors typically initiate Smad signaling through C-terminal phosphorylation of R-Smads. We therefore examined effects of shear stress magnitude (1-30 dynes/cm^2^) on Smad 2/3 C-terminal phosphorylation, over a time course of 12 hours. We also examined C-terminal phosphorylation of Smad 1/5/8 for comparison. In both cases, Smad phosphorylation peaked around 2-4 hours after the onset of flow and then decreased to a plateau that was above baseline by 12 hours (Figure 2A-B). However, unlike nuclear translocation, there was no decrease in phosphorylation at higher shear stress for Smad 2/3. We also examined phosphorylation of the linker region, which marks R-Smads for degradation. Although oscillatory flow (OSS) showed a trend toward higher linker phosphorylation and lower total Smad 2/3 levels, there were no statistically significant effects (Figure 2C). Thus, while flow induces C-terminal phosphorylation of Smad 2/3, it does not show the same biphasic dependence on FSS magnitude as nuclear translocation. We conclude that flow initiates Smad 2/3 activation via phosphorylation but that inhibition of nuclear localization and gene induction at high FSS is a separate effect.

**Figure 2.**
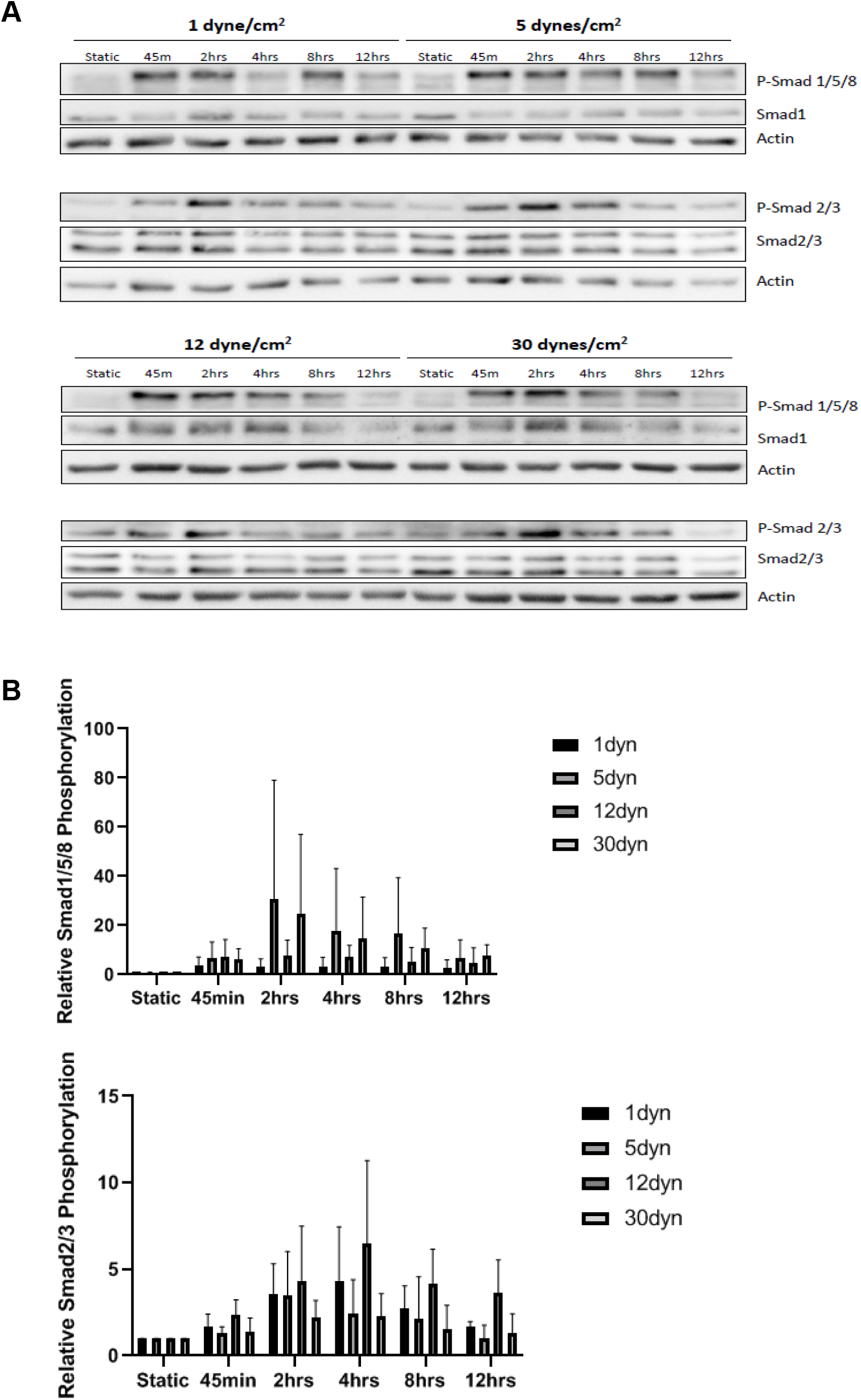
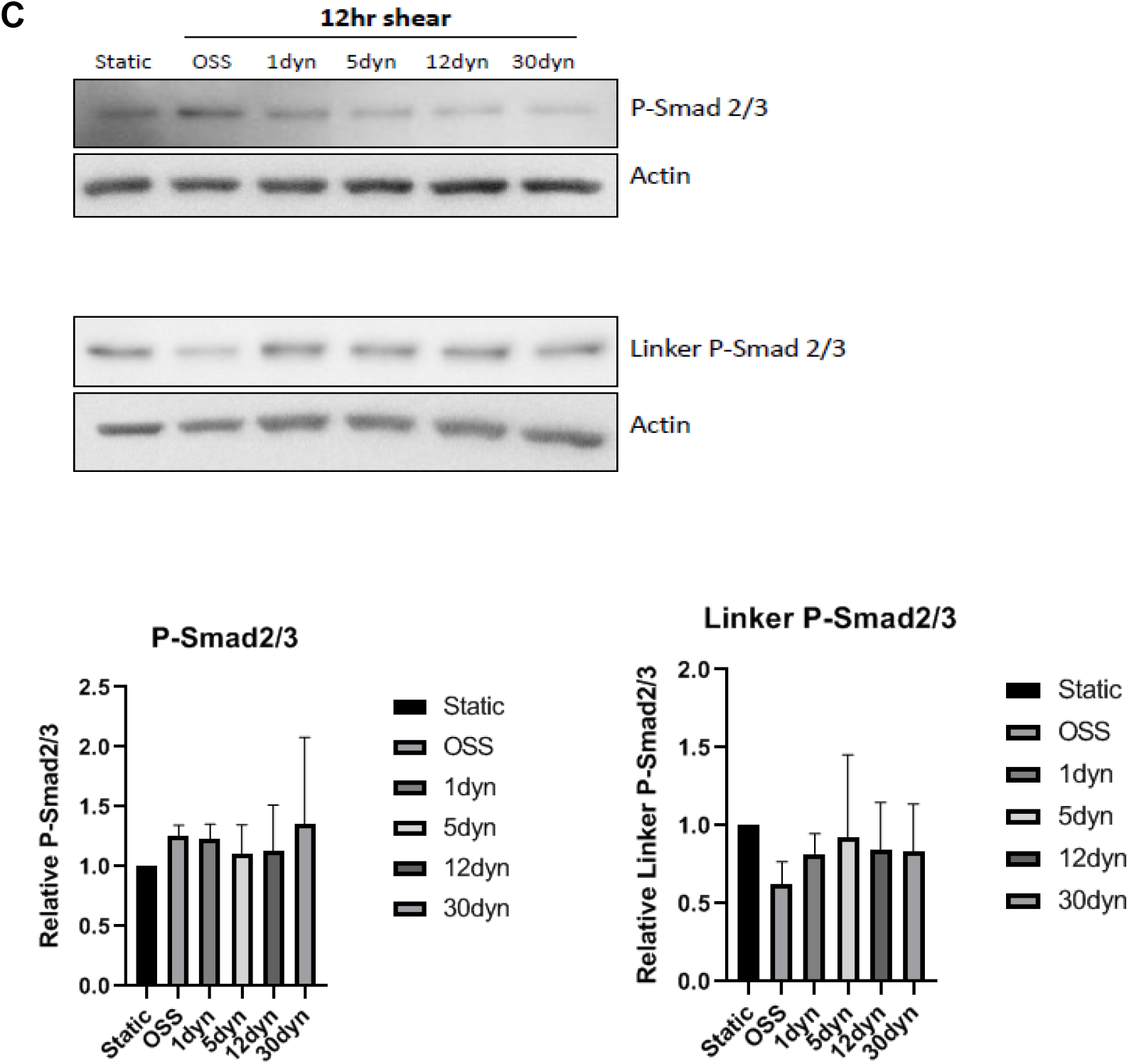
Smad 2/3 C-terminal and linker phosphorylation. **A:** HUVECs were exposed to LSS at the indicated magnitudes for 12 hours. C-terminal phosphorylation of Smad 1/5/8 and Smad 2/3 was assessed by Western blotting of cell lysates. **B:** Quantification of Smad 1/5/8 and Smad 2/3 phosphorylation results from A. Values are means ± SEM, n=3. **C:** Linker phosphorylation levels of Smad 2/3 measured by Western blotting of cell lysates and results quantified as above. Values are means ± SEM, n=3. ^*^ p<0.05, ^**^ p<0.01

### Identification of receptors

Previous work from our lab identified Alk1 and Endoglin receptors as key mediators of flow-induced Smad 1/5/8 activation [11]. To determine which receptors mediate flow activation of Smad 2/3, we first considered Alk5, the major Type I receptor for TGFβ-induced Smad 2/3 activation. In parallel, we examined Alk1 and Endoglin, and effects on Smad 1/5. As expected, siRNA-mediated depletion of Alk5 in HUVECs inhibited C-terminal Smad 2/3 phosphorylation (Figure 3A). Re-expression of siRNA-resistant Alk5 rescued this effect (Figure 3B). While depletion of Alk1 or Endoglin blocked flow activation of Smad 1/5, there were no detectable changes in Smad 2/3 phosphorylation. We also considered Neuropilin-1 (Nrp1) based on recent results reporting an interaction with Alk1 and 5, and its association with another receptor, VEGFR2, that is implicated in flow signaling [17]. Depleting Nrp1 reduced Smad 2/3 C-terminal phosphorylation under activation by flow, which was rescued by re-expression (Figure 3C) but had no effect on Smad1/5 activation by flow. It also had no effect on BMP9-induced Smad 2/3 phosphorylation under static conditions (Figure 3D). These results demonstrate that Alk5 and Nrp1 are the receptors required for flow activation of Smad 2/3.

**Figure 3.**
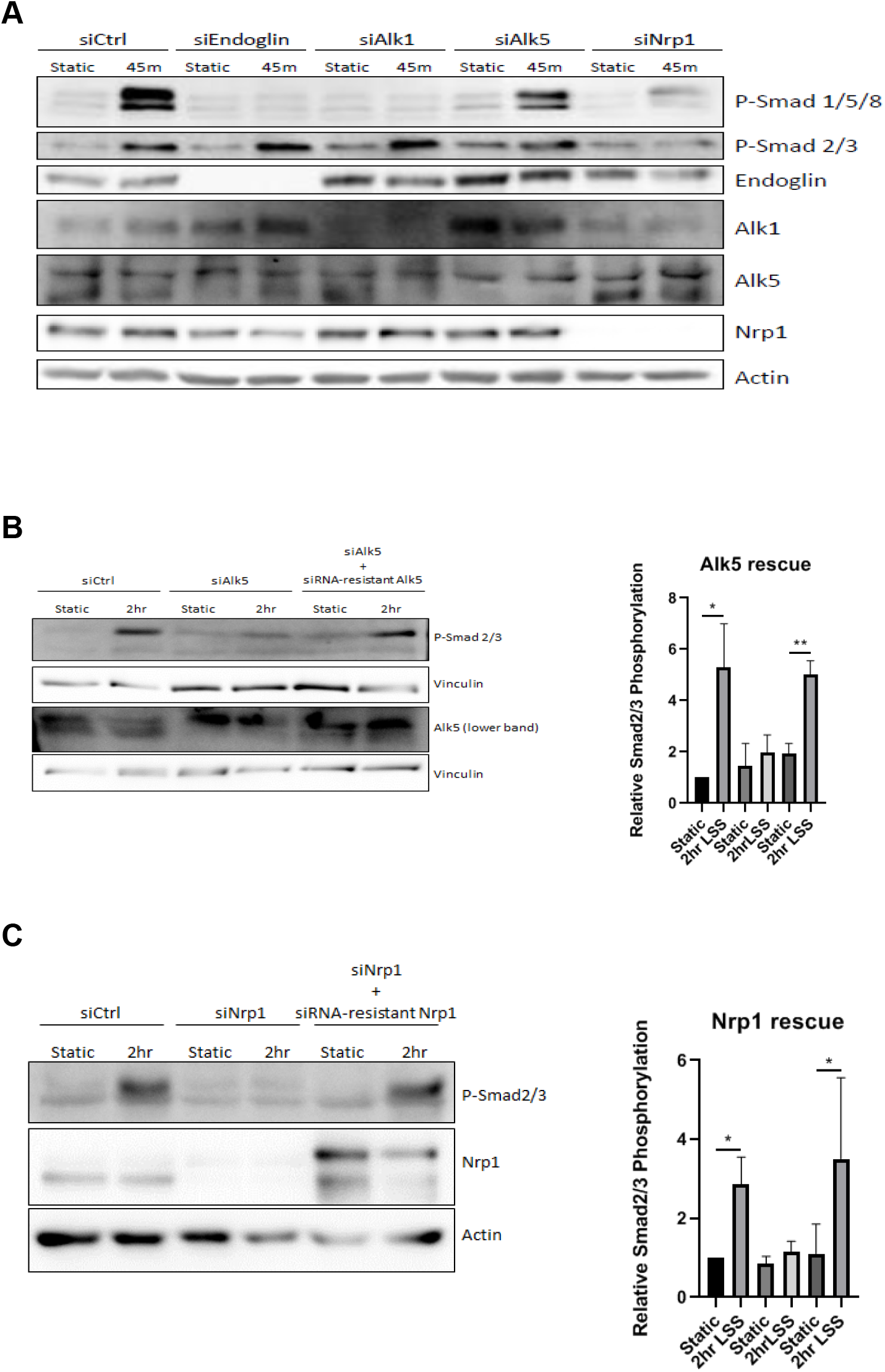
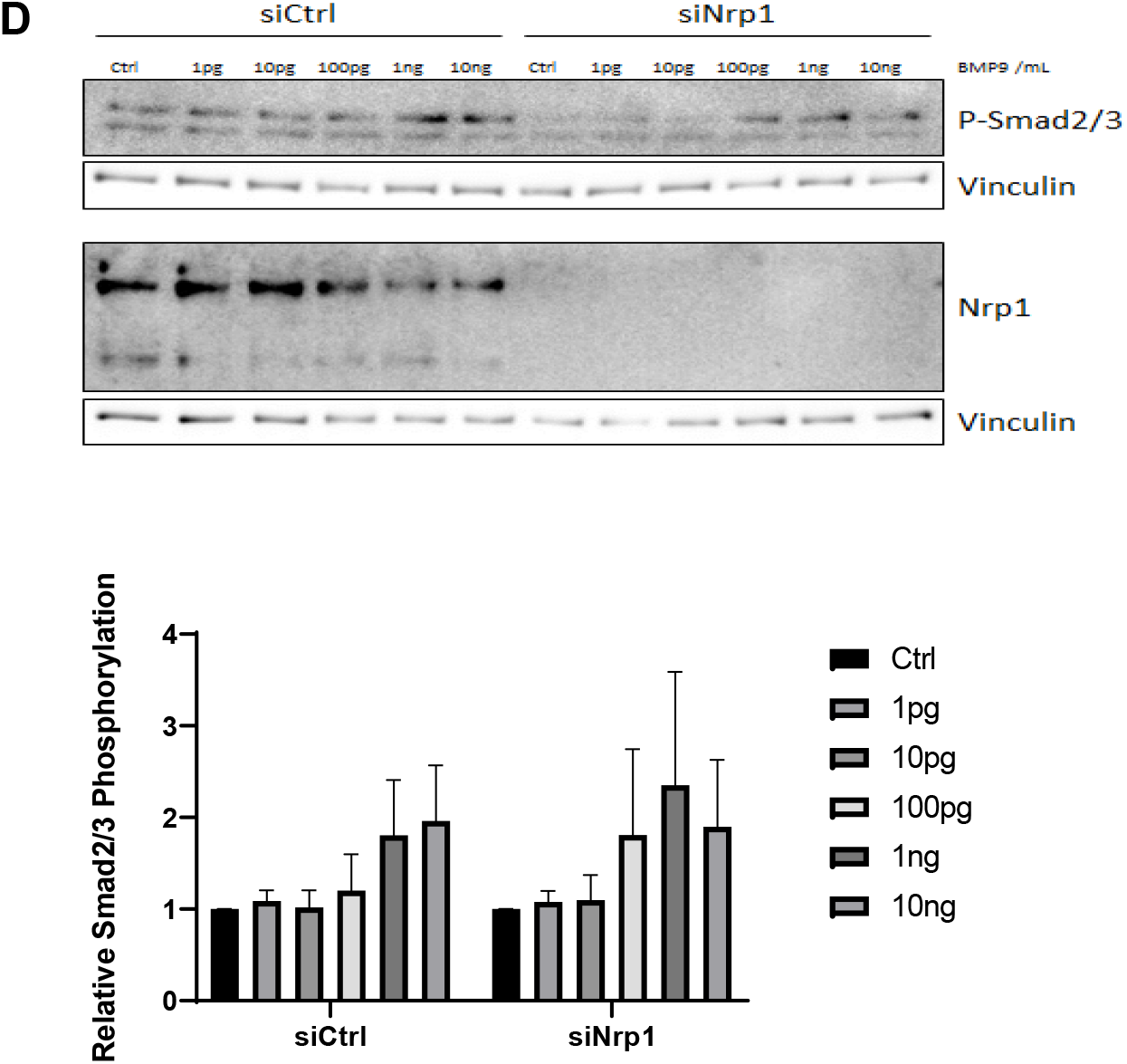
Identification of receptors for flow-induced Smad 2/3 activation. **A:** ECs were subject to knockdown of the indicated receptors and flow activation of Smads assayed by Western blotting. **B:** Alk5 knockdowns were rescued by re-expression of siRNA-resistant Alk5 and flow activation of Smad 2/3 assayed. For quantification, Values are means ± SEM, n=3. **C:** Nrp1 knockdown was rescued by re-expression of siRNA-resistant Nrp1 and flow activation of Smad2/3 assayed. For quantification, Values are means ± SEM, n=3. **D:** ECs after Nrp1 depletion were stimulated with the indicated concentration of BMP9 in the absence of shear stress and Smad 2/3 activation assayed by Western blotting. Quantified values are means ± SEM, n=3. ^*^ p<0.05, ^**^ p<0.01

### Identification of ligands

We next set out to identify the ligands involved in LSS-mediated Smad 2/3 activation. We first tested TGFβ2, which is the major Alk5 ligand that stimulates Smad 2/3 activation. HUVECs in serum-free medium with 0.2% BSA were treated with TGFβ2 at concentrations ranging from 1pg/mL to 10ng/mL, without or with FSS. However, shear stress had no effect on the dose-response curve for TGFβ2-induced Smad 2/3 phosphorylation (Figure 4A). Flow also showed a trend toward weak suppression of the maximal Smad 2/3 activation but this effect did not reach statistical significance. Given that Smad 1/5 activation by BMP9 and BMP10 were enhanced by flow, we next tested these ligands. Flow induced a leftward shift in the dose response curve for BMP9-induced Smad 2/3 activation by about 10-fold, similar to the effect on Smad 1/5 activation (Figure 4B). To our surprise, flow did not affect the response to BMP10 (Figure 4C). To confirm these findings, we exposed HUVECs in medium with 2% serum to LSS without or with BMP-blocking antibodies. Anti-BMP9 reduced flow-induced Smad 1/5/8 and Smad 2/3 phosphorylation while anti-BMP10 only reduced flow-induced Smad 1/5/8 phosphorylation (Figure 4D). We conclude that the FSS enhancement of Smad 2/3 activation is specific to BMP9 but not related ligands.

**Figure 4:**
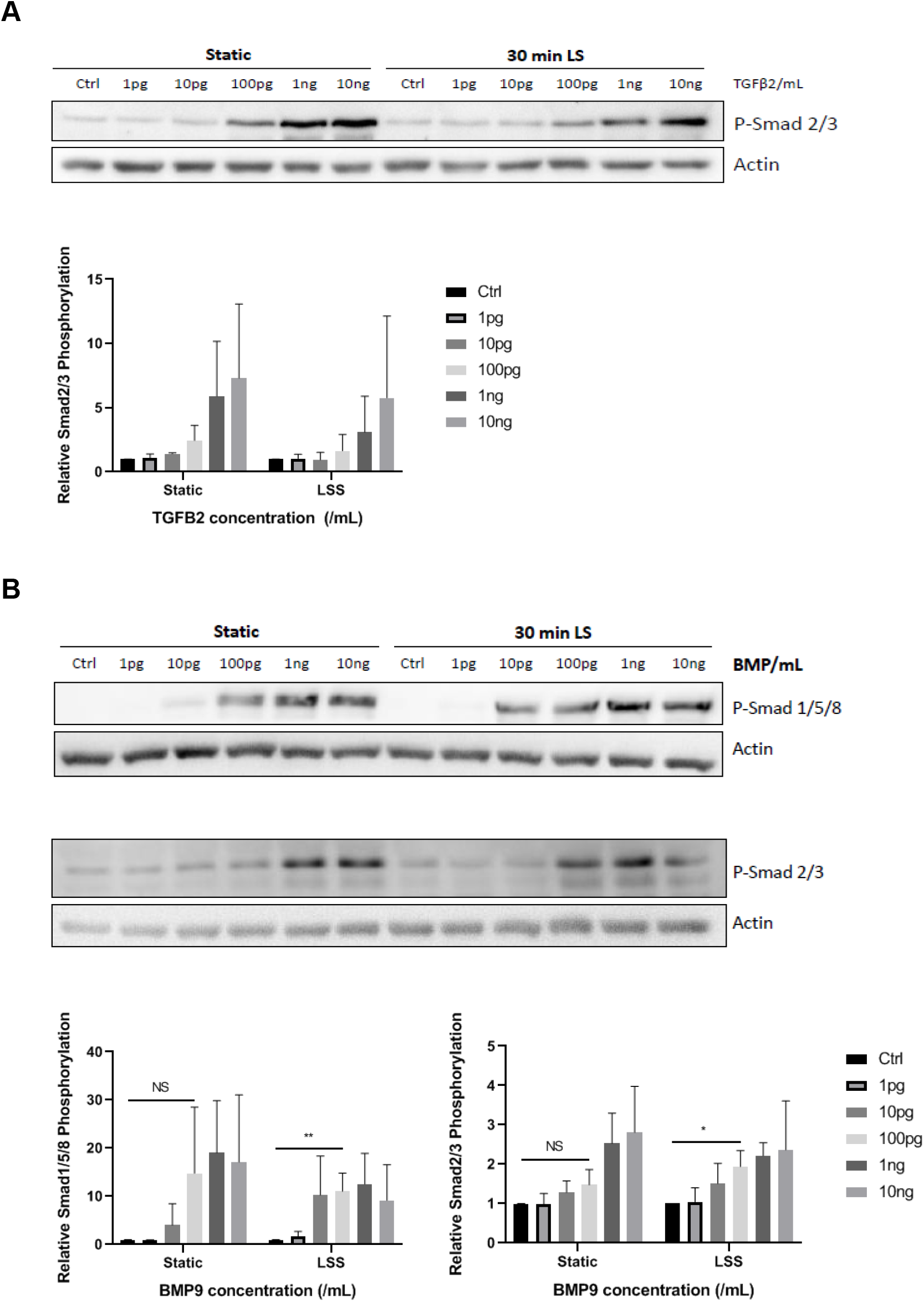
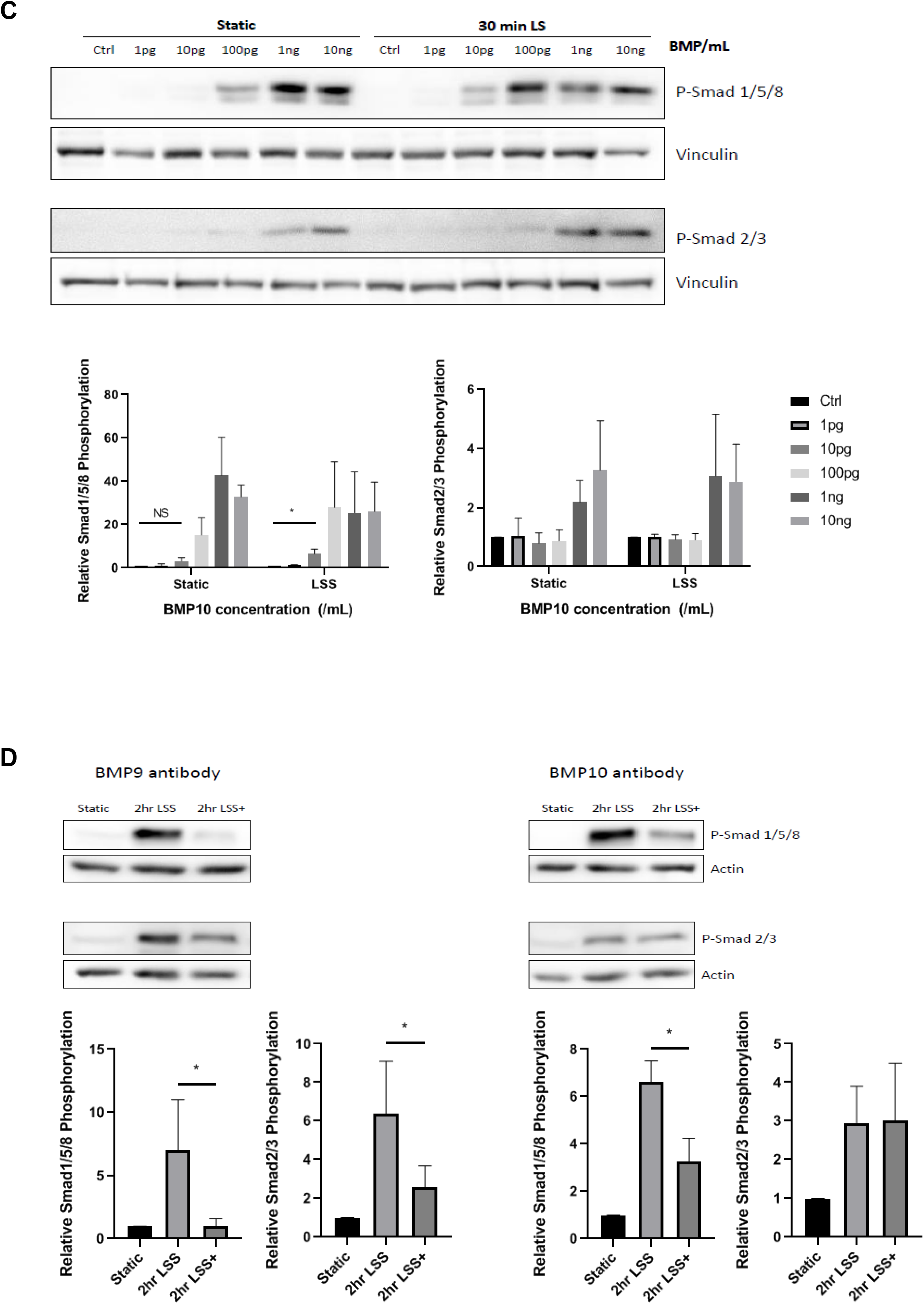
Identification of ligands for flow-induced Smad 2/3 activation. ECs were stimulated by addition of the indicated concentrations of ligands in the presence or absence of LSS for 30 min, and Smad 2/3 phosphorylation assayed by Western blotting **A:** TGFβ2. Results were quantified from 3 experiments. Values are means ± SEM. **B:** BMP9. Results were quantified from 3 experiments. Values are means ± SEM. **C:** BMP10. Results were quantified from 3 experiments. Values are means ± SEM. **D:** HUVECs in 2% serum were exposed to LSS for 2h without (2hr LSS) or with (2hr LSS+) BMP-blocking antibodies to BMP9 or 10 as indicated. Samples were then assayed for Smad phosphorylation and results quantified from 4 experiments. Values are means ± SEM. ^*^ p<0.05, ^**^ p<0.01

### Low flow-induced inward remodeling in vivo

We next considered whether this pathway controls low flow-mediated inward remodeling in vivo. We used a model in which ligation of two of the branches off the right carotid (Figure 5A) reduces flow by around 75-80% and inward remodeling to ~50% of the initial diameter or circumference. There is also a slight compensatory increase in flow in the contralateral carotid but outward remodeling is limited to about 10%. Right carotid ligation in WT mice triggered an increase in endothelial P-Smad 2 staining in the common carotid 5 mm upstream from the closure, demonstrating Smad 2 activation by low flow in vivo (Figure 5B). To test whether Smad 2 activation is required for artery remodeling, we used Cdh5:Cre, Alk5^fl/fl^ mice for tamoxifen-inducible, EC-specific deletion of Alk5. Eight-week old mice were injected with tamoxifen for five days and subject to surgery one week later. Two weeks after surgery, arteries were fixed, harvested, and stained with H&E. Lumen circumference measurements showed that in WT mice, the right carotid artery circumference decreased by 18% (p<0.0039), which corresponds to a 33% decrease in area. In Alk5 ECKO mice, there was no statistically significant decrease (Figure 5C). Altogether, these data demonstrate that Alk5-Smad 2 signaling contributes to low flow-induced inward artery remodeling.

**Figure 5:**
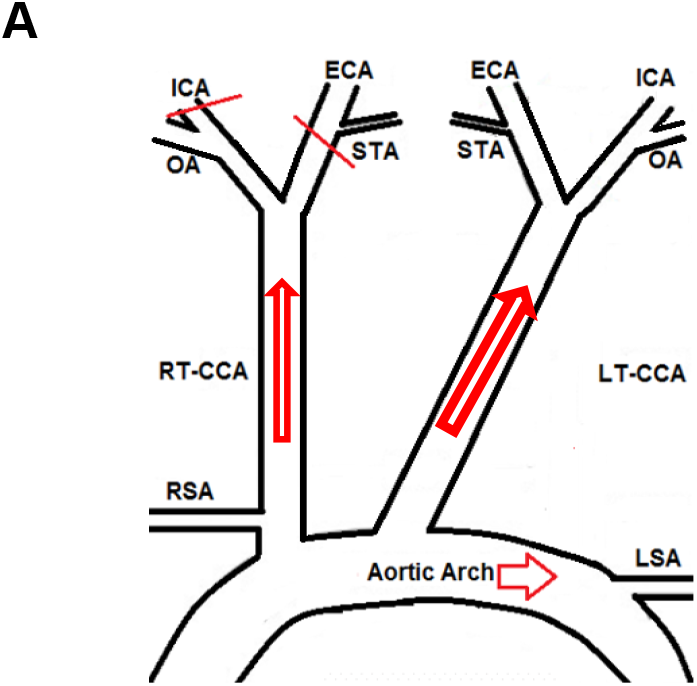
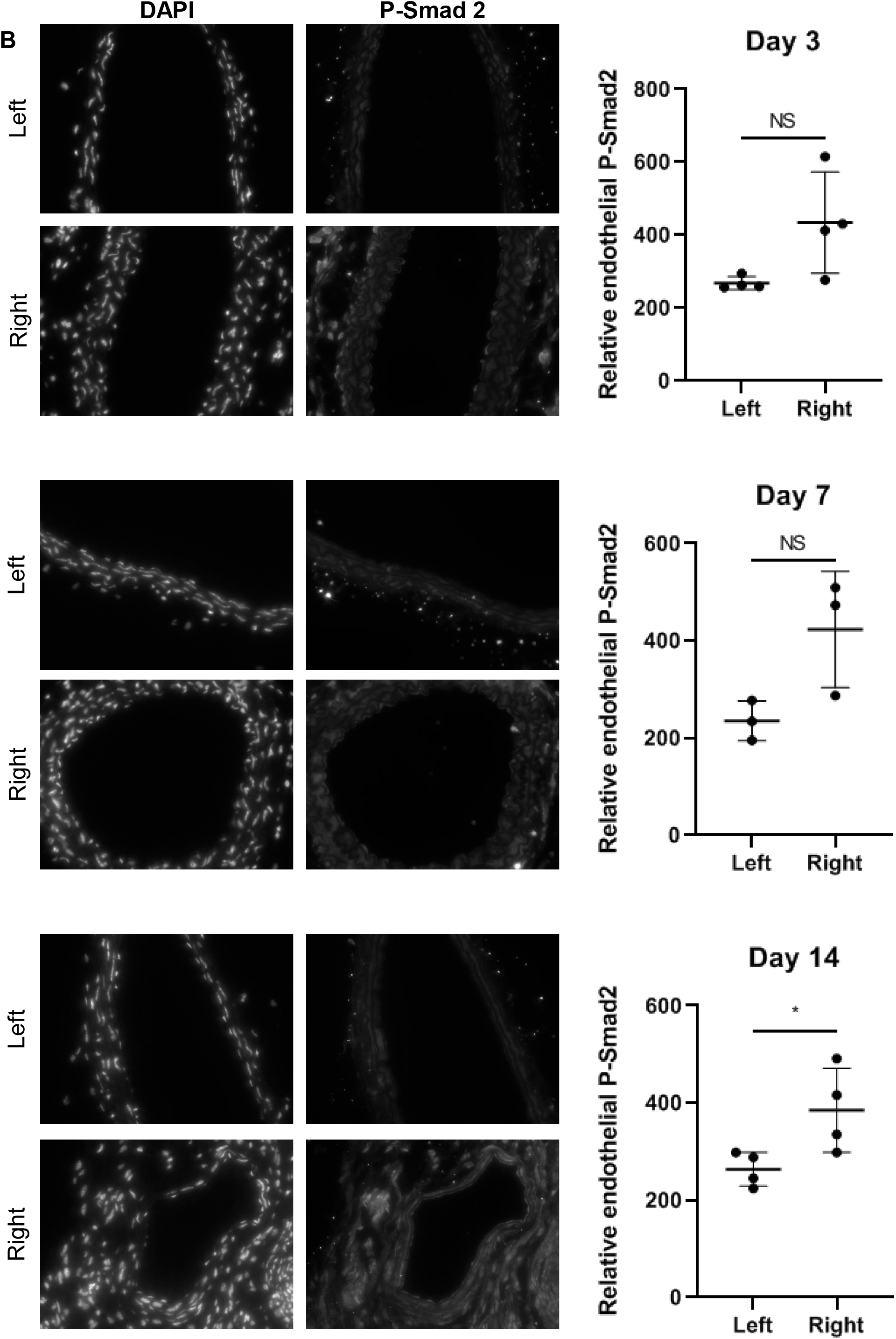
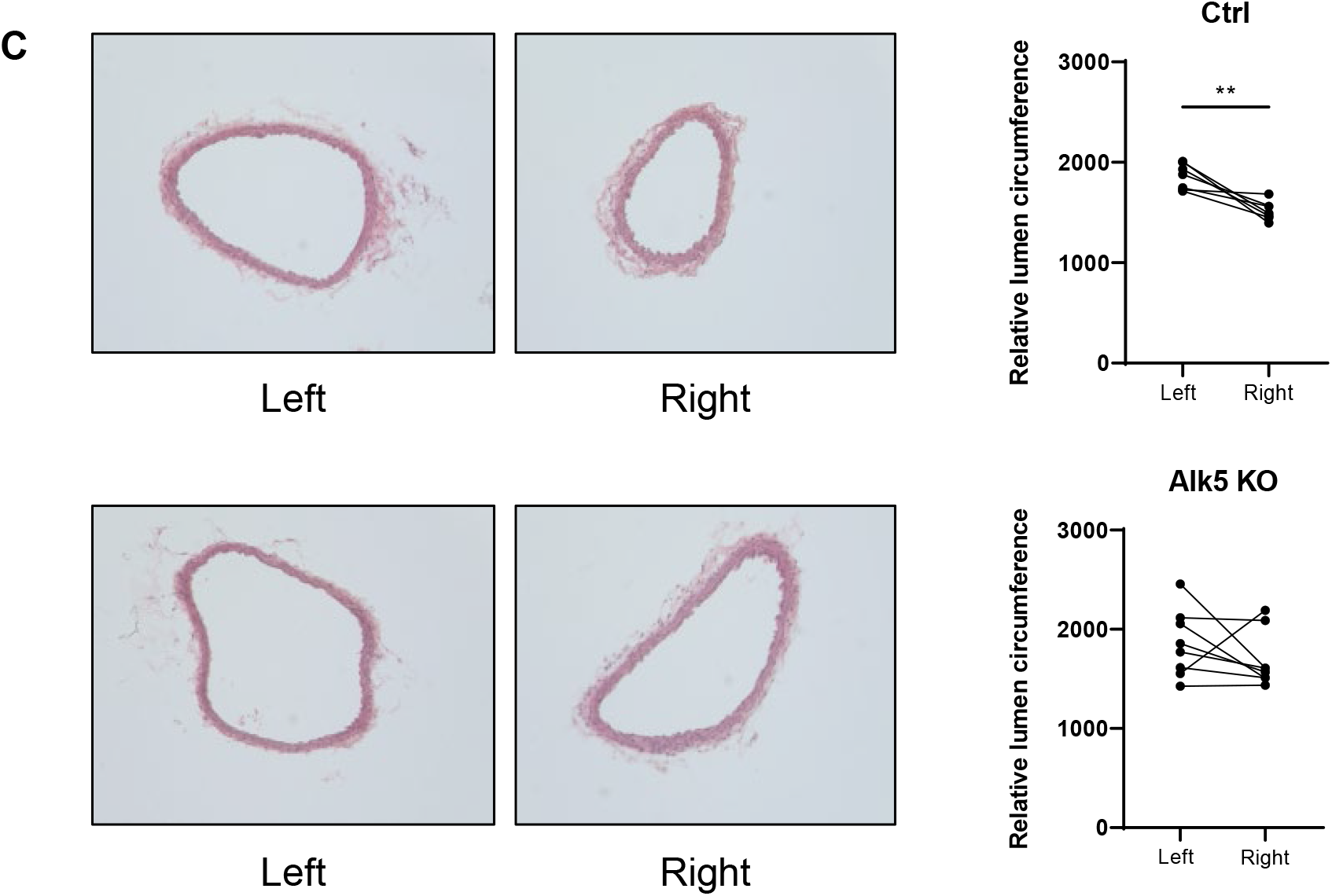
Low flow-induced inward remodeling in vivo. **A:** Schematic of carotid ligation: Three branches of the right common carotid artery (RCA), the external carotid artery (ECA), internal carotid artery (ICA), and the superior thyroid artery (STA) were ligated, leaving the occipital artery (OA) open. **B:** Carotid ligation as illustrated in Fig. 5A was performed on the right side in WT mice. Four mice were sacrificed for each time point post-surgery. Three sections of each carotid artery were stained with P-Smad 2 antibody and quantified. **C:** Mice with or without EC-specific deletion of Alk5 were subject to right carotid ligation and vessel circumference assessed at 2 weeks.^*^ p<0.05, ^**^ p<0.01

## Discussion

In this study, we demonstrate that FSS activates Smad 2/3 and induces nuclear translocation and gene expression maximally at low shear levels, which promotes low flow-induced inward artery remodeling. This pathway has several unconventional features that raise interesting new questions. First, FSS triggers C-terminal phosphorylation of Smad 2/3 with a monotonic dose-dependence, that is, phosphorylation increases from 1 dyne/cm^2^ and then plateaus at higher forces (e.g., 5 to 30 dynes/cm^2^). However, high shear (12 dynes/cm^2^ and above) reduces nuclear translocation of the phosphorylated Smads and target gene expression. The reduced nuclear localization or cytoplasmic retention at high shear thus confers maximal signaling at low levels of FSS, consistent with inward remodeling. The molecular mechanism by which high shear inhibits nuclear translocation will be an important direction for future efforts.

Smad 2/3 activation by flow is due to a shift in the dose-response for BMP9, a known ligand for Alk5. BMP9 is synthesized mainly in the liver and circulates at high levels in early embryos but persists at lower levels through adulthood [18]. Surprisingly, although BMP10 has 65% sequence similarity and is functionally similar in terms of Smad 1/5/8 activation, it does not share in the ability to induce flow-dependent Smad 2/3 activation. Our data also identify Alk5 and Nrp1 as the key receptors in the flow-dependent activation of Smad 2/3. Neither receptor has a role in flow or BMP9 activation of Smad 1/5. While Alk5 is well known as an activator of Smad 2/3, the involvement of Nrp1 is surprising and molecular mechanism unclear. Nrp1 was observed to interact with and modulate the activation state of Alk5 and Alk1, as well as functions of other receptors including VEGF receptors and integrins [17]. This function of Nrp1 in Smad 2/3 activation by flow could conceivably be linked to its interaction with VEGF receptors, which also participate in flow signaling [19, 20]. Indeed, changing expression of VEGF receptors shifts the FSS set point between different types of endothelial cells [6]. Future work will be required to elucidate the molecular mechanisms by which Nrp1 mediates flow activation of this pathway.

These results are likely relevant to Smad 2/3 activation by low/oscillatory shear stress in atherosclerosis-prone regions of arteries [13]. In that setting, flow activation of 2/3 likely contributes to lesion formation at the initiation phase. Smad 2/3 activation further increases in atherosclerosis at later stages, which involves interactions with inflammatory factors and TGFβ, thereby promoting endothelial-mesenchymal transition and more severe inflammation and pathological vessel wall remodeling. Thus, in this setting, Smad 2/3 appears to be part of a positive feedback circuit that promotes disease progression. Severe disease is often accompanied by decreased lumen diameter that reduces flow, ultimately closing or restricting arteries even without apparent rupture or thrombosis. The current finding that Smad 2/3 promotes inward artery remodeling may thus be relevant to clinical progression in severe artery disease. Future work to elucidate molecular mechanisms and clinical consequences is thus an important direction for future research

